# Prioritization of chemical scaffolds using the TDR Targets database: an integrative workflow for *Trypanosoma cruzi* drug discovery

**DOI:** 10.64898/2026.03.02.709050

**Authors:** Lionel Urán Landaburu, Mercedes Didier Garnham, Emir Salas Sarduy, Fernán Agüero

## Abstract

Chagas disease, caused by the parasite *Trypanosoma cruzi*, faces a critical innovation gap in drug development, with current treatments hindered by toxicity and limited efficacy. To address this, we implemented an integrative chemogenomic workflow using the TDR Targets database to prioritize drug candidates. To prioritize repurposing candidates for *T. cruzi*, we designed a query to retrieve compounds active against validated targets in other organisms, provided an orthologous gene exists in *T. cruzi* and the compound has no recorded activity against trypanosomatids and their associations predicted by the TDR Targets multilayer network. On those associations we applied sequential filters based on metabolic relevance, and commercial availability via the MolPort API obtaining a focused set of 378 high-priority compounds.

A central feature of this workflow was the partitioning of these compounds into 16 distinct chemical libraries, each defined by unique scaffolds such as benzamidines, sulfonamides, and azoles. For experimental validation, we manually curated two of these libraries, containing piperazine and nitro derivatives. From the 21 compounds acquired for *in vitro* testing against *T. cruzi* in intracellular models of infection, 7 demonstrated selective trypanocidal activity, with two lead hits achieving submicromolar EC_50_ values.

Crucially, while our experimental focus was on these two series, the remaining 14 curated libraries, representing a broad range of chemical space and putative target associations, which are fully available for public exploration and further biological assaying. These results demonstrate the efficiency of our prioritization pipeline and provide the scientific community with a pre-filtered, commercially accessible resource to accelerate the discovery of new leads for Chagas disease.

**Author summary:** Chagas disease, caused by the parasite *Trypanosoma cruzi*, is a neglected tropical disease with limited treatment options. To accelerate drug discovery, we developed an integrative computational workflow using the TDR Targets database as a starting point to prioritize compounds for repurposing. We retrieved compounds active against validated targets in other organisms, ensuring they had no recorded activity against trypanosomatids and that their associations were predicted by the TDR Targets multilayer network. Based on this work we provide a 12 chemical libraries with 378 prioritized chemical scaffolds for furhter experimental validation. In this work we focused on two of these sublibraries for experimental validation, and from 21 compounds tested, 7 showed selective activity against the parasite. The remaining curated chemical libraries, represent a broad range of chemical space and putative target associations, and are fully available for exploration and further biological assaying. This study demonstrates the efficiency of our prioritization pipeline and provides a valuable resource to accelerate the discovery of new leads for Chagas disease.

## Introduction

The World Health Organization (WHO) has identified 20 Neglected Tropical Diseases (NTDs), which have historically affected impoverished populations across Africa, Asia, and the Americas [1–3]. Among them, Chagas disease represents a major social, economic and public health burden in Latin America and is recognized by the World Health Organization (WHO) as one of the most prevalent NTDs [4]. Chagas disease is caused by the protozoan parasite *Trypanosoma cruzi*. It is endemic throughout the American continent, affecting approximately 8 million people [5], with an increasing number of cases reported in North America [6]. Clinically, the disease manifests in two distinct phases: a short, mostly asymptomatic acute phase and a decades-long chronic phase, associated with severe complications such as megacolon and potentially fatal cardiomyopathies [7].

Currently, there is no available vaccine for Chagas disease and existing treatment options rely on Nifurtimox (NFX) and Benznidazole (BNZ), two drugs with limited efficacy in chronic infections and considerable toxicity [8]. Thus, the development of new, effective, and safer therapeutic alternatives remains a pressing need. However, the limited commercial incentive for pharmaceutical companies to invest in treatments for NTDs, primarily due to low expected financial returns, has resulted in a striking innovation gap: only 1% of new drugs approved between 1977 and 2002 target these diseases, and none of them for Chagas disease [9, 10]. From 2000 to 2011, neglected diseases received just 4% of all new products and 1% of new chemical entities [11].

Given the high costs and extended timelines required for *de novo* drug discovery, such strategies have historically been limited for NTDs [12]. Addressing this gap requires the systematic integration of heterogeneous biological and chemical data, a field known as chemogenomics. While global drug repurposing efforts have identified promising compounds for NTDs [13], the available pathogen-specific data remains critically sparse compared to the vast resources for model organisms [2]. This scarcity hence requires data from well-characterized model organisms, particularly where core cellular processes are evolutionarily conserved. Dedicated bioinformatics strategies are essential for this cross-species integration. Resources like the TDR Targets database [14–17] enable the ranking of potential drug targets and the identification of bioactive compounds by systematically integrating genomic data from pathogens with functional annotations and chemical bioactivity data from model systems [2, 3]. This interconnected architecture is particularly suited for NTDs, as it provides a robust framework to leverage bioactivity data from extensively studied organisms to guide drug discovery in pathogens that typically lack high-volume, experimental chemogenomic datasets.

The TDR Targets database integrates a chemogenomics network that is built on three primary entity data types [16, 17]: bioactive chemical compounds, protein targets from pathogen and model organism proteomes, and functional annotation nodes (e.g., Pfam domains [18], OrthoMCL ortholog groups [19], KEGG pathways [20]). Distinct layers encode specific relationship types: a compound-compound layer based on structural similarity (Tanimoto coefficient), a protein-annotation affiliation layer that groups targets based on shared annotations, and crucially, a compound-protein bioactivity layer populated by manually curated and literature-derived interactions [16]. This interconnected structure formalizes the chemogenomic space, allowing for graph-based analyses and inference. A key analytical output derived from this network is the Network Druggability Score (NDS), a metric designed to prioritize protein targets that can be associated with known compounds with higher inference confidence. The NDS algorithm evaluates the local topology surrounding each target node and assigns relevance weights to shared annotation nodes based on the statistical over-representation of known druggable (compound-linked) proteins within those functional categories [17]. A target’s NDS, which ranges from 0 to 1, is thus a weighted composite score reflecting the density and confidence of connections to bioactive compounds via informative functional annotations. A high NDS indicates not only that a target resides in a network region enriched with successful pharmacological interactions, but more importantly, that it can be prioritized with greater confidence for compound assignment. This provides a data-driven, multi-evidence proxy for druggability that extends beyond simple sequence similarity.

While these computational frameworks are robust, a significant challenge remains in translating an abstract chemogenomic graph (often containing millions of bioactivity records) into tangible, testable hypotheses. In this study, we explicitly leverage the multilayer framework of TDR Targets as the foundational step in a novel drug discovery pipeline for *T. cruzi*.

When the multilayered network was first generated [16], the NDS score was used to perform a prioritization of the best ten targets within the ‘TriTryp’ kinetoplastid parasites (*Trypanosoma cruzi, T. brucei*, and *Leishmania major*) [16]. Furthermore, other groups have used the TDR Targets-network based engine to guide prioritization of targets and compounds [21–32].

The TDR Targets platform underwent a significant expansion with the release of TDR Targets version 6, which integrated 22 new genomes and extensive updates to chemical and bioactivity data [17]. This version enabled a new ranking of complete proteomes, including *T. cruzi*, based on the NDS. This also led to the definition of 5 Druggability Groups (DGs) or Tiers for targets, with targets classified in group 5 (DG5) representing the highest scores and greatest confidence, and DG1 the lowest. While the foundational work demonstrated the model’s robustness by identifying established targets [17], these prioritizations served primarily as a proof-of-concept and remained without prospective experimental validation.

In this work, we move beyond *in silico* prioritization to experimental validation. Using the TDR Targets chemogenomic framework, we selected high-confidence *T. cruzi* targets and retrieved all associated bioactive compounds—roughly 180,000 molecules. Through sequential filtering based on metabolic relevance and commercial availability, we narrowed these to 378 candidates representing diverse chemical scaffolds. From 2 selected libraries, we tested 21 compounds *in vitro* and identified 7 with selective trypanocidal activity and low cytotoxicity. This demonstrates that network-based predictions can deliver novel, biologically active compounds, providing a generalizable strategy for drug discovery in neglected diseases.

## Materials and methods

### Construction of the Screening Libraries

The construction of the libraries was carried out in two stages. First, several datasets were collected from multiple sources, with TDR Targets being one of the most relevant. The complete list of datasets is available in Table 1.

**Table 1.** List of datasets collected for generation of the screening libraries.

| Source | Dataset | Label | Approx. Size | Type<sup>a</sup> |
| --- | --- | --- | --- | --- |
| TDR Targets | NDPI<sup>b</sup> | Compounds obtained by transforming a list of targets into a list of compounds | 180,023 | (C) |
| TDR Targets | Tested | Compounds tested (active, inactive, inconclusive) against trypanosomatids | 6619 | (C) |
| DrugBank | Drugbank | FDA-approved drugs, marketed and withdrawn | 11,172 | (C) |
| Molport | Available | Commercial availability | $> 5 \times 10^6$ | (C) |
| Various<sup>c</sup> | NDPI<sup>b</sup> | Manually curated high-throughput screening hits against *T. cruzi* not yet included in TDR Targets | 36 | (C) |
| TDR Targets | TDR-Targets-D | *T. cruzi* proteins with druggability group 4 | 327 | (P) |
| TDR Targets | TDR-Targets-EA | *T. cruzi* proteins involved in energy and amino acid metabolism | 44 | (P) |

<sup>a</sup> Dataset type: (C) compound library; (P) protein target set.
<sup>b</sup> network-derived putative interactions
<sup>c</sup> Manually compiled dataset from phenotypic screenings [33,34]

Once collected, the datasets were sequentially combined in order to reduce the number of entities to a manageable size. Initially, all records from the full list of network-derived putative interactions whose compounds were also found in the tested compounds list were removed. The resulting dataset was intersected with the commercially available compounds dataset (available), eliminating all records involving compounds that could not be commercially sourced, thereby generating a new dataset. This was then combined with the DrugBank dataset in an attempt to obtain a quick list of repositionable drugs; however, no interesting candidates were identified.

Putative targets were also used to reduce the size of the original dataset. To do this, the dataset resulting from the previous operations was intersected with a curated list of genes of interest, both highly druggable (TDR-Targets-D) and presumably involved in energy and amino acid metabolism (TDR-Targets-EA) [35].

The compound library was subdivided into subsets of molecules containing a particular functional group or core scaffold. The representative scaffolds are shown in Table 2. For experimental validation, two of these libraries were selected for compound refinement, resulting in a library of 31 piperazine derivatives and another containing 16 nitro-containing compounds. From this total of 47 compounds, 21 were commercially acquired for *in vitro* assays.

**Table 2.**
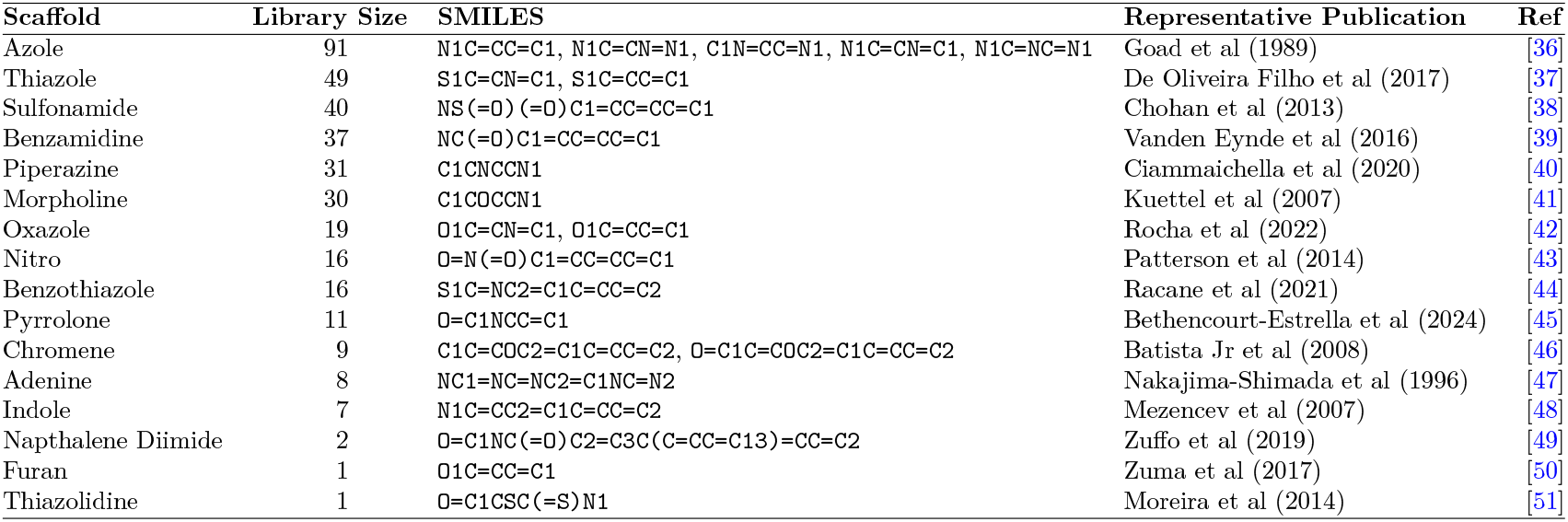
Scaffold-based compound libraries.

| Scaffold | Library Size | SMILES | Representative Publication | Ref |
| --- | --- | --- | --- | --- |
| Azole | 91 | N1C=CC=C1, N1C=CN=N1, C1N=CC=N1, N1C=CN=C1, N1C=NC=N1 | Goad et al (1989) | [36] |
| Thiazole | 49 | S1C=CN=C1, S1C=CC=C1 | De Oliveira Filho et al (2017) | [37] |
| Sulfonamide | 40 | NS(=O)(=O)C1=CC=CC=C1 | Chohan et al (2013) | [38] |
| Benzamidine | 37 | NC(=O)C1=CC=CC=C1 | Vanden Eynde et al (2016) | [39] |
| Piperazine | 31 | C1CNCCN1 | Ciammaichella et al (2020) | [40] |
| Morpholine | 30 | C1COCCN1 | Kuettel et al (2007) | [41] |
| Oxazole | 19 | O1C=CN=C1, O1C=CC=C1 | Rocha et al (2022) | [42] |
| Nitro | 16 | O=N(=O)C1=CC=CC=C1 | Patterson et al (2014) | [43] |
| Benzothiazole | 16 | S1C=NC2=C1C=CC=C2 | Racane et al (2021) | [44] |
| Pyrrolone | 11 | O=C1NCC=C1 | Bethencourt-Estrella et al (2024) | [45] |
| Chromene | 9 | C1C=COC2=C1C=CC=C2, O=C1C=COC2=C1C=CC=C2 | Batista Jr et al (2008) | [46] |
| Adenine | 8 | NC1=NC=NC2=C1NC=N2 | Nakajima-Shimada et al (1996) | [47] |
| Indole | 7 | N1C=CC2=C1C=CC=C2 | Mezencev et al (2007) | [48] |
| Napthalene Diimide | 2 | O=C1NC(=O)C2=C3C(C=CC=C13)=CC=C2 | Zuffo et al (2019) | [49] |
| Furan | 1 | O1C=CC=C1 | Zuma et al (2017) | [50] |
| Thiazolidine | 1 | O=C1CSC(=S)N1 | Moreira et al (2014) | [51] |

The Python code used to construct and subdivide the libraries, together with the resulting scaffold-specific subsets, is available on GitHub as a Jupyter notebook (see Data Availability).

### Chemical compounds and reagents

Benznidazole (Abarax) powder was a gift from Laboratorios Elea, Argentina. The stock solution was prepared in dimethyl sulfoxide (DMSO) at 20 mM. Other investigational compounds were sourced from the following suppliers through Molport (Riga, Latvia) and Hit2Lead (ChemBridge Corporation, CA, USA) (Table S2). These compounds were solids and some required solubilization in a small amount of DMSO before dilution in aqueous media.

### Cytotoxicity determination

To determine cytotoxicity, a resazurin (RZ) assay was performed using the same compound concentrations on non-infected Vero cell cultures [52]. Cells were seeded in 96-well plates at a density of 5 *×* 10^3^ cells/well and cultured for 48 hours. They were then carefully washed with PBS, fed with fresh RPMI medium supplemented with each compound (20 *µ*M), and incubated for 72 hours. Afterward, 10 *µ*L of freshly prepared 10X (440 *µ*M) RZ solution was added to each well. The assay included BNZ (20 *µ*M) and DMSO (0.5%) as controls, along with an untreated cell control and a no-cell control to establish the maximum and minimum reagent reduction readings. In addition, a 100% reduced RZ control (1X RZ solution sterilized by autoclaving in RPMI medium) was included on the plate as a reference to determine when to take the final reduction reading: absorbance was measured at 600 nm every hour until the untreated cell control reached 100% of the reduced RZ control reading. The final measurement was taken after 7 hours of incubation, recording absorbance at both 600 nm and 570 nm. All measurements were performed using a FilterMax F5 Multimode Microplate Reader (Molecular Devices, CA, USA). Compounds and controls were tested in duplicate. For the initial screening, cytotoxicity was measured as a function of production of resorufin (reduction product of resazurin), using Equation 1, where F is fluorescence measurement and *µ*C+ represents the mean of vehicle controls.

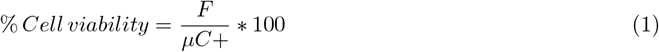

Compounds maintaining ≥ 35% Vero cell viability at the screening concentration were considered for subsequent CC_50_ evaluation based on their *T. cruzi* growth inhibition profiles. For CC_50_ determination, dose–response cytotoxicity assays were performed under the same experimental conditions described above, using at least eight two-fold serial dilutions of each compound. The highest concentrations assayed were 100 *µ*M for CID-5, 50 *µ*M for CID-6 and CID-19, 20 *µ*M for CID-7, and 7.5 *µ*M for CID-21. All conditions were assayed in duplicate or triplicated. Data were imported into GraphPad Prism version 8.0.1 (GraphPad Software LLC) and analyzed using nonlinear regression to generate log(dose)-response curves with a variable slope. CC_50_values, defined as the concentration causing a 50% reduction in cell viability, were calculated independently for each compound.

### Parasite Culture

*T. cruzi* Tulahuen strain parasites constitutively expressing *E. coli β*-galactosidase (Tul *β*-gal) were a generous gift of Frederick Buckner (University of Washington, USA). Initial stocks of infectious trypomastigotes were obtained through serial passages in CF1 mice. At the peak of parasitemia, transgenic Tul-*β*-gal parasites were obtained from anticoagulated blood of infected mice. For purification, on the same day as the mouse bleeding, the blood sample was diluted with 1 or 2 volumes of PBS and centrifuged at 300 g for 5 minutes; it was then incubated at 37 °C for 40 minutes to promote swimming of the trypomastigotes. This centrifugation and incubation (swimming) step was repeated a second time. After 40 minutes, the supernatant was again collected and centrifuged at 4,500 g for 7-10 minutes to pellet the swimming trypomastigotes. The parasite pellet was washed repeatedly with RPMI medium, with centrifugation between each wash. Before the final wash, an aliquot was taken to count and assess parasite viability.

For all infection assays, cell-derived Tul *β*-gal trypomastigotes were propagated in Vero cells maintained at 37 ^◦^C with 5% CO_2_ in a humidified incubator. Cells were cultured in minimum essential medium (MEM; Gibco Life Technologies) supplemented with 10% fetal bovine serum, 100 U/mL penicillin, and 10 *µ*g/mL streptomycin. Trypomastigotes were collected from culture supernatants 96 hours post-infection by centrifugation at 5,000 x g for 10 minutes. Parasites were routinely maintained in Vero cells for up to 40 passages without detectable loss of *β*-galactosidase activity. Thereafter, new cultures were initiated from cryopreserved bloodstream trypomastigotes.

### Trypanocidal activity determination

To determine the trypanocidal activity of the investigational compounds, Tul *β*-gal parasites were used. The enzymatic activity of the *β*-galactosidase (*β*-gal) reporter enzyme was used as a proxy for parasite growth and measured via a colorimetric end-point assay using chlorophenol red-*β*-D-galactopyranoside (CPRG) as substrate [53]. For this assay, 5 *×* 10^4^ trypomastigotes per well were used to infect Vero cells previously seeded (24 h before) in a 96-well plate at a density of 5 *×* 10^3^ cells per well. After 24 hours of infection, cultures were gently washed with PBS to remove free trypomastigotes and then incubated in phenol red-free RPMI medium (Gibco Cat No. 11835030) supplemented with the compound (20 *µ*M). Benznidazole (BNZ), also at 20 *µ*M, was used as a positive control, along with DMSO (0.5%) as a negative/vehicle control, and untreated infected and uninfected controls to establish maximum and minimum *β*-gal activity readings, respectively.

After incubating the culture with the compound (20 *µ*M, 2 *µ*M, or 0.2 *µ*M) for 96 hours, 100 *µ*L of a freshly prepared solution containing 1% nonyl phenoxypolyethoxylethanol (NP-40) and 100 *µ*M CPRG (Roche Cat No. 10884308001) was added to each well, reaching final concentrations of 0.5% NP-40 and 50 *µ*M CPRG per well. The plates were then incubated for 4 hours at 37 °C, protected from light throughout the incubation. Finally, *β*-gal activity was measured by reading absorbance at 595 nm using a FilterMax F5 Multimode Microplate Reader (Molecular Devices). All compounds were tested in duplicate.

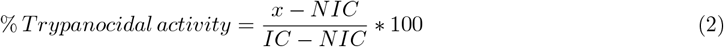

The inhibition of parasite growth was calculated using Equation 2, where x represents the absorbance measurement of the sample, IC is the absorbance of the infected control (maximum parasite growth), and NIC is the absorbance of the non-infected control (minimum parasite growth). Compounds exhibiting ≥ 60% *T. cruzi* growth inhibition and 35% Vero cell viability at the screening concentration were classified as direct hits and considered for subsequent evaluation in EC_50_ dose-response assays. For compounds exhibiting 60% *T. cruzi* growth inhibition but ≤ 35% Vero cell viability at the screening concentration, the trypanocidal assay was repeated under identical experimental conditions, but concentrations were reduced to 2 *µ*M and 0.2 *µ*M.

All EC_50_ dose-response experiments were performed in 96-well plates using the same experimental setup described above. Compounds were tested starting at concentrations up to 100 *µ*M for CID-5, 50 *µ*M for CID-6 and CID-19, 20 *µ*M for CID-7, and 7.5 *µ*M for CID-21 using nine serial 1:2 dilutions. All experimental conditions were performed in duplicate. Trypanocidal activity was calculated using Equation 2 and data was imported into GraphPad Prism version 8.0.1 (GraphPad Software LLC) for analysis. EC_50_ values were calculated independently for each compound using a nonlinear regression model to generate log(dose)-response curves with a variable slope. The selectivity index (SI) was calculated for each compound as the ratio of CC_50_ to EC_50_.

### Beta-galactosidase inhibition assayxs

Selected compounds were tested for inhibition of *β*-galactosidase activity to eliminate potential false positives, ie., compounds able to mimic true trypanocidal drugs by interfering with the reporter enzymatic assay. An extract of transgenic *T. cruzi* Tul *β*-gal parasites was used as the source of *β*-galactosidase. Transgenic parasites expressing *β*-galactosidase were obtained from the culture medium of infected cells, as previously described. The supernatant from a cell culture flask was centrifuged at 5,700 x g for 10 minutes and the pellet was washed twice with PBS. The resulting pellet was resuspended in 100 *µ*l of PBS, supplemented with 0.5% NP-40 and 1 *µ*M E-64 protease inhibitor, and incubated for 1 hour at 37 °C. After incubation, the parasite lysate was sonicated in four 5-second cycles at 5% amplitude, followed by centrifugation at 16,200 x g for 30 minutes at 4 °C. The supernatant was collected, supplemented with 20% glycerol, and stored at −80 °C until use.

Compounds were preincubated for 30 min at 37 °C with a *β*-galactosidase–containing lysate (1:2000 dilution) in activity buffer (PBS supplemented with 2% RPMI, 0.25% DMSO, and 0.5% NP-40, pH 7.4). The CPRG substrate was then added to a final concentration of 50 *µ*M. Data acquisition and analysis were performed as described above. Residual *β*-galactosidase activity was calculated for each condition using Equation 1 and used to generate dose–response curves. Isopropyl *β*-D-1-thiogalactopyranoside (IPTG, 0-5 mM) served as a *β*-galactosidase inhibition control.

## Results

### Data Exploration and Filtering Strategy

To identify compounds with potential for repurposing against *T. cruzi*, a query was designed to retrieve molecules known to be active against a validated target in another organism, provided that an orthologous gene exists in *T. cruzi* and that the compound has no recorded activity against *T. cruzi, T. brucei*, or *L. major*. This strategy enables the prioritization of candidates that may act on *T. cruzi* through conserved biological targets but have not yet been evaluated in that specific context. With this query we selected a subset of 327 genes. From these protein coding genes, we then extracted the entire set of bioactive compounds associated with them by TDR Targets (approximately 180,000).

To bridge the gap between associations derived from the chemogenomic network and experimental feasibility, the network derived putative interactions set was subjected to a multi-stage filtering process using a set of Python scripts (available in a Jupyter Notebook). First, a novelty filter was applied to exclude compounds already present in DrugBank [54] or previously tested against trypanosomatids, leaving a refined set of 178,144 molecules. Recognizing that commercial accessibility is a critical bottleneck in neglected disease research, we programmatically validated the entire set via the MolPort API, finding that only 15% (29,880 compounds) had a similar compound available for purchase at the time of this assessment and that 28,802 were an exact match with the ones prioritized in this work. This intersection of selected genes and compounds narrowed the scope to 44 core metabolic genes associated with 4,041 commercially available compounds, serving as the definitive starting point for structural refinement of these set of compounds.

Then 44 genes were prioritized due to their involvement in energy and amino acid metabolism. This *ad hoc* filtering step was based on evidence indicating that carbon metabolism is notably enhanced in cells infected with amastigotes [55], as well as the role of certain amino acids (and intermediate metabolites) in parasite survival during infection [56].

The final stage of the pipeline focused on structural clustering and outlier removal to ensure the selection of high-quality chemical leads. Using MACCS keys and an all-vs-all Tanimoto similarity matrix, we implemented a hierarchical single-linkage clustering approach. A distance (or height) threshold of 0.8 was selected to generate coherent microclusters while preventing the conflation of distinct scaffolds. To mitigate pharmacological promiscuity, we performed a noise removal step by discarding compounds linked to more than 10 different target families, a strategy analogous to PAINS analysis [57, 58]. This systematic curation yielded a final set of 378 high-priority compounds associated with 42 genes. The list of genes and the number of compounds associated with each one is shown in Table S1. Figure 1 shows the order in which the datasets were combined and the number of features obtained in each case.

**Fig 1.**
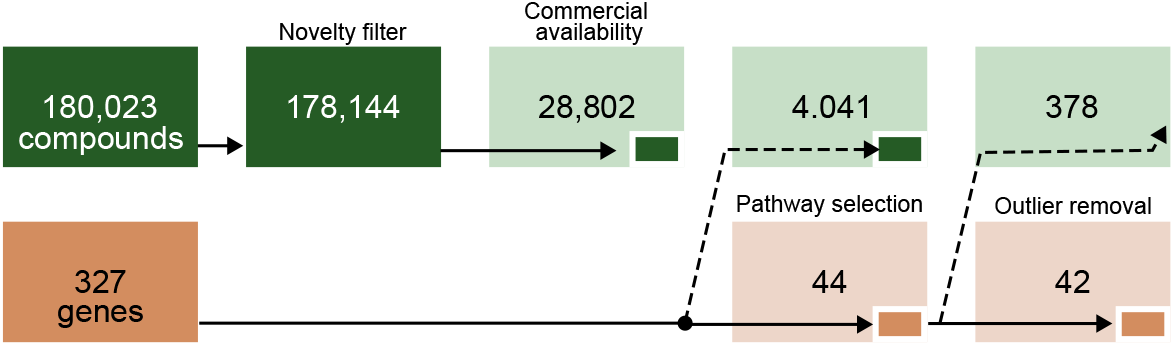
Compound and target gene selection workflow. The initial dataset of 180,023 compounds and 327 target genes was sequentially filtered by novelty and commercial availability, resulting in 28,802 compounds. Pathway-based selection reduced the dataset to 4,041 compounds and 44 genes, followed by outlier removal yielding a final set of 378 compounds and 42 genes.

### Construction of Screening Libraries

Based on the cleaned clustered data, a substructure search was performed for sets of functional groups reported in the literature as potentially trypanocidal (see Methods). The compound candidates were partitioned into 16 distinct scaffold-based libraries. The complete list of scaffolds can be found in Table 2. Using this information, we partitioned our dataset into focused libraries: each library was characterized by containing compounds featuring one or more of these functional groups. The libraries were not exclusive (a molecule may belong to one or more libraries). This sub-grouping aimed to facilitate the final exploration of the libraries. All the libraries are available through Github (see Data availability section).

For experimental validation we selected the nitro and piperazine chemical libraries guided by two distinct yet complementary rationales. Nitro-containing compounds were prioritized due to their well-documented history as reactive (class-specific) pharmacophores with potent anti-*Trypanosoma cruzi* activity [43]. In contrast, piperazines were selected not for intrinsic reactivity, but for their versatility as structural scaffolds and linkers. The piperazine scaffold provides conformational flexibility and multiple points for chemical diversification, enabling the presentation of a wide range of appended functional groups with the potential to engage diverse biological targets (generalistic). Together, the inclusion of these two libraries reflects a strategy to incorporate chemical sets with different origins and mechanistic rationales: one based on known parasite-specific chemical reactivity and the other on scaffold-driven structural versatility and exploratory potential. From these two libraries we selected 21 compounds to test *in vitro* (Figure 2 ). The selection reflected mainly short term availability and price affordability at the time of purchase.

**Fig 2.**
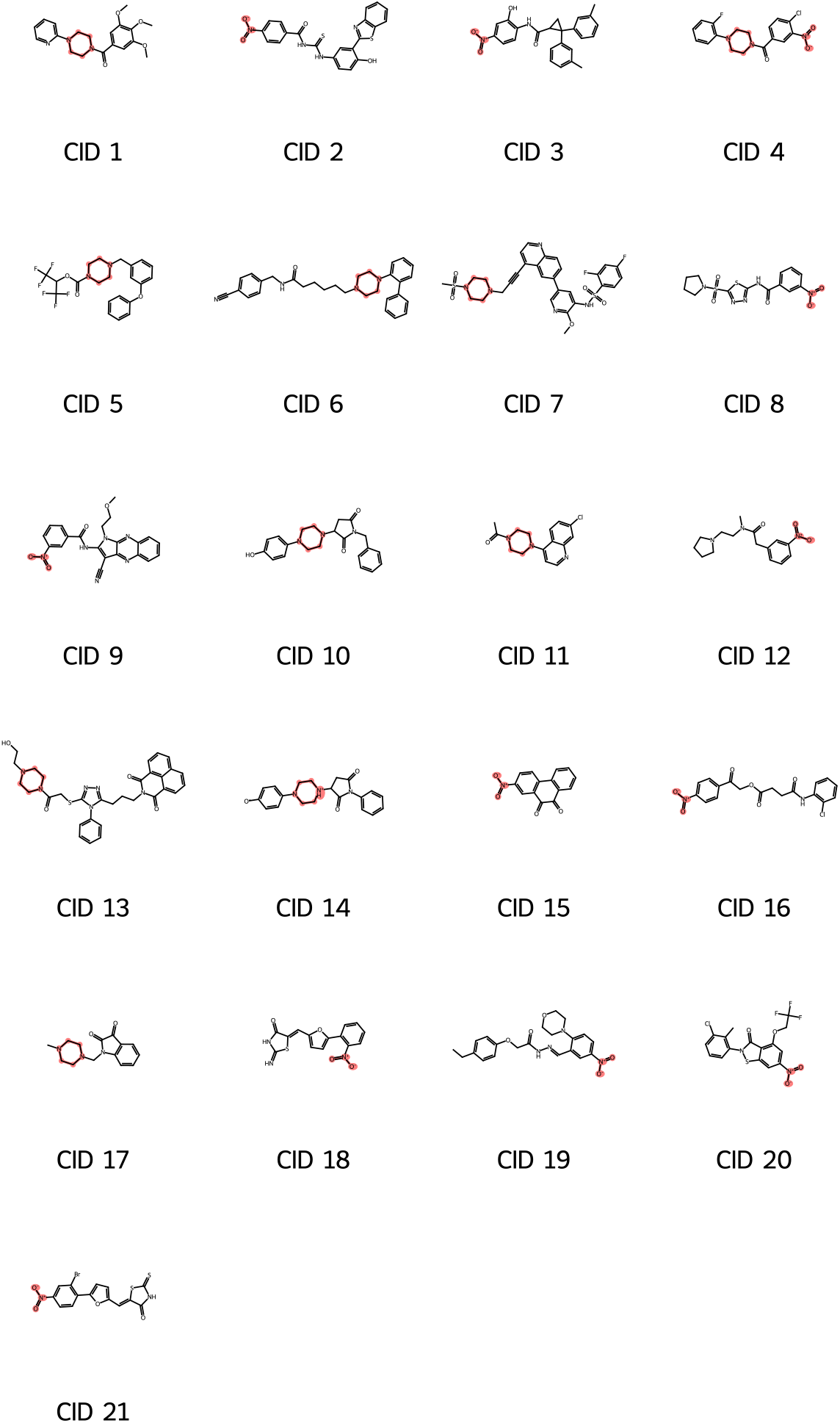
Selected compounds for testing. The piperazine and nitro scaffolds, which define the core chemical series explored in this work, are highlighted.

### Experimental Validation: Primary Screening

In a primary screening, the 21 selected compounds were tested *in vitro* to assess their cytotoxicity and trypanocidal activity at a fixed concentration of 20 *µ*M (see Methods). Results from primary screening are summarized in Figure 3.

**Fig 3.**
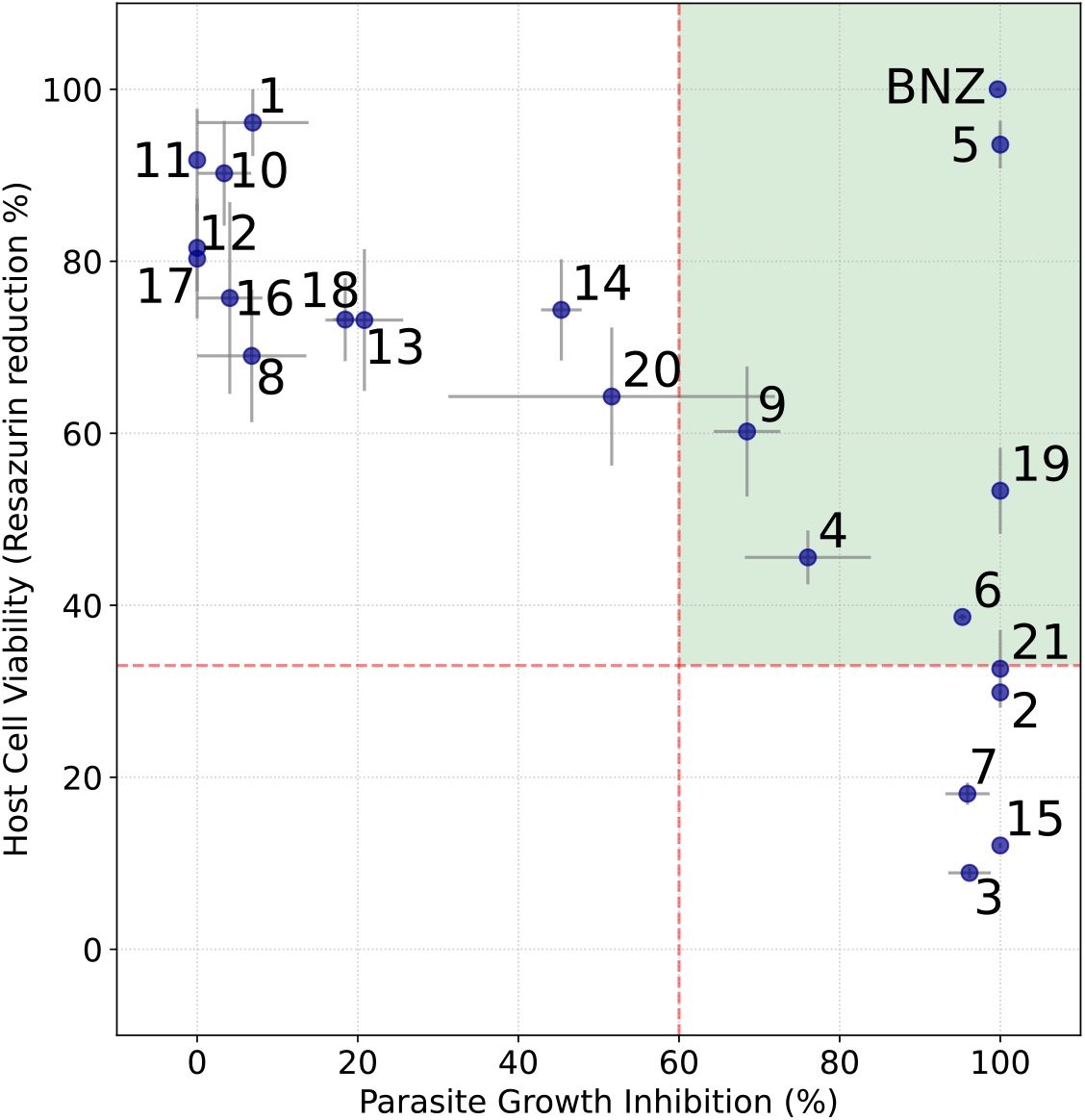
Trypanocidal activity vs. cytotoxicity plot from primary screening. Each compound was tested in duplicated at a fixed dose of 20 *µ*M. Results from the *β*-galactosidase activity and resazurin reduction assays are shown for each compound. The horizontal axis represents the percentage of *T. cruzi* amastigote growth inhibition based on *β*-gal activity measured with the CPRG substrate. The vertical axis shows cell viability as a function of the percentage of resazurin reduction at a fixed time point: the higher the percentage, the higher the cellular metabolism, and thus the lower the compound’s cytotoxicity. Horizontal and vertical bars for each point represent measurement errors, expressed as the standard deviation between replicates and their average. For this assay, thresholds were set at 35% for cytotoxicity and 60% for growth inhibition. Compounds of interest appear in the upper right quadrant (high inhibition of parasite growth, high cellular viability) and are labeled as ‘CID-number’, along with BNZ (Benznidazole), which was used as a positive control. Compounds in the lower right quadrant were also selected for retesting at lower concentrations. Arrows point to compounds that gave satisfactory results when retested at reduced concentrations. Molecules in the left quadrants were considered negative.

While 11 compounds showed only moderate or no antiparasitic activity (inhibition% < 60%), other 11 compounds, including the reference drug BNZ, significantly inhibited parasite growth at 20 *µ*M on the primary screening. Of them, CID-2, CID-3, CID-5, CID-6, CID-7, CID-15, CID-19, and CID-21, completely arrested parasite growth (% inhibition> 90%) at this concentration, standing out as the most promising candidates. Contrary to their uniform antiparasitic activity, these compounds displayed highly disparate toxicity on Vero cells. Five compounds (CID-2, CID-3, CID-7, CID-15, and CID-21) were highly toxic (cell viability % < 35%) at 20 *µ*M, and three (CID-5, CID-6, and CID-19) exhibited moderate to low cytotoxicity.

Of note, CID-5 showed the most favourable activity/cytotoxicity profile among the investigated candidates, with almost maximal anti-*T. cruzi* activity and negligible cytotoxicity on host cells.

To explore for differential effects on parasite and cell viability at other concentration ranges, the five compounds showing the highest cytotoxicity were re-tested at lower doses of 2 *µ*M and 0.2 *µ*M (Figure S1). CID-2 failed to reproduce the bioactivity observed in the previous screening round and CID-15 showed similar effects on both parasites and Vero cells, indicating indiscriminate toxicity. Thus, both compounds were discarded. On the contrary, CID-3, CID-7, and CID-21 exhibited highly selective bioactivity profiles at 2 *µ*M. At this concentration, these compounds inhibited parasite growth almost completely (Inhibition % > 95%) while demonstrating high cell viability (70%-85%), indicative of acceptable selectivity ratios. All the compounds showed only negligible bioactivity at the lowest concentration tested (0.2 *µ*M).

In summary, after primary screening, 8 compounds (CID-3, CID-4, CID-5, CID-6, CID-7, CID-9, CID-19, and CID-21) remained as viable candidates, as they showed reproducible and selective antiparasitic activity (Figure 3 ). The top five hits CID-5, CID-6, CID-7, CID-19, and CID-21 were prioritized for further dose-response analysis in secondary screening.

### Dose-response analysis and enzymatic triaging of top hits

Dose-response assays were performed to estimate half-maximal effective (EC_50_) and cytotoxic (CC_50_) concentrations of each compound under standardized conditions. As expected from primary screening, the analysis revealed marked differences among compounds in both antiparasitic potency and selectivity. The most potent and selective molecule, CID-7, exhibited a submicromolar EC_50_ (5.15 *×* 10^−7^ M) together with the highest selectivity index (SI = 29.26), indicating a favorable therapeutic window. CID-5 and CID-19 also showed antiparasitic potencies in the low micromolar range, with EC_50_ values of 7.50 *×* 10^−6^ M and 5.84 *×* 10^−6^ M, respectively, and moderate selectivity. Despite its strong potency against *T. cruzi* (EC_50_ = 1.85 *×* 10^−6^ M), CID-21 displayed only a modest selectivity index due to a CC_50_ value in the two-digits micromolar range. Among the selected hits, CID-6 was the least promising candidate. Although no signs of cytotoxicity were detected in dose-response analysis, this compound showed the lowest antiparasitic potency (EC_50_ = 2.68 *×* 10^−5^ M) of the series, resulting in a poor selectivity index (SI > 1.87). Figure 4 summarizes dose–response curves and calculated parameters for these compounds.

**Fig 4.**
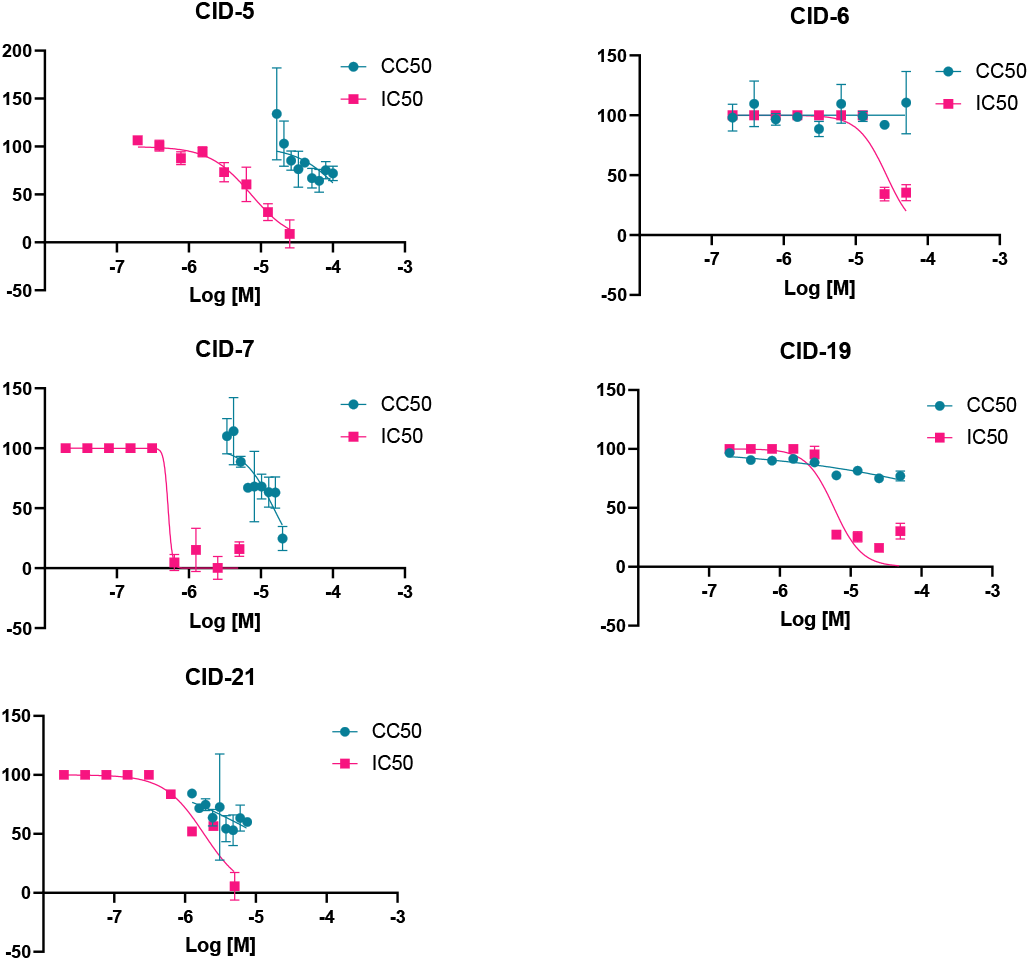
Dose-response curves for the estimation of CC_50_ and EC_50_ in confirmatory screening. Selected hits from primary screening were re-tested across a range of concentrations to generate dose-response curves, from which EC_50_ and CC_50_ were estimated. Curves for the antiparasitic and cytotoxic activity of each compound are indicated in fuchsia and cyan, respectively. Each point represents the mean and SD of 3 three technical replicates.

Finally, the investigated compounds were tested in an enzymatic triage to discard candidates able to interfere with the reporter methodology (i.e., *β*-galactosidase inhibitors or highly coloured compounds) used in the screening assay. In the range of concentrations tested, none of the investigated compounds showed detectable *β*-galactosidase inhibition (Figure S2), indicating that all the hits genuinely reduced parasite burden in culture. The estimated parameters and the chemical structures of the compounds assayed in secondary screening are summarized in Table 3. Overall, our results validate CID-5, CID-7, CID-19, and CID-21 as *bona fide* hits with selective anti-*T*.*cruzi* activity with potential as starting point for further optimization and confirm the libraries generated from TDR Targets as enriched sources of attractive candidates.

**Table 3.**
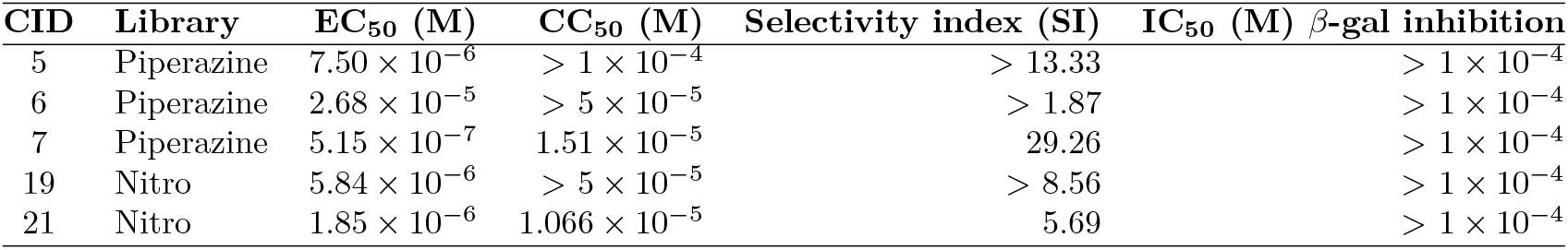
Compound activity data against *T. cruzi*.

## Discussion

Despite decades of intense research, Chagas disease remains the most important parasitic disease in the Americas, with the only two drugs in clinical use displaying suboptimal efficacy and safety profiles. To cope with the urgent need for new and safer therapeutic options for this disease, in this work we developed an integrative workflow to prioritize chemical scaffolds with potential activity against *T. cruzi*. Using the TDR Targets Database, we first applied an *ad hoc* filtering strategy to identify molecules with confirmed activity against orthologous targets in other organisms but no recorded activity against *T. cruzi* or related kinetoplastids (*T. brucei* and *L. major*). This generated a set of *T. cruzi* targets from which we extracted all compounds associated with direct neighbors in the TDR Targets chemogenomic network (see voting scheme prioritization in Berenstein *et al*. [16]). Then we enriched the list with high potential, actionable candidates by using an *ad hoc* filtering strategy. As a result, 16 scaffold-specific libraries, encompassing more than 4,000 drug-like compounds, were assembled and made available as a public resource that can be interrogated in phenotypic screening to identify novel hits. Our experimental validation with two unrelated sublibraries (21 compounds) permitted us to validate at least four novel candidates (two from each library) combining micromolar-to-submicromolar potencies with suitable selectivity indices.

The TDR Targets database is a valuable resource that can be used in various ways. From its initial conception and through successive updates, the intention has been to create a flexible and autonomous web platform, providing tools to manipulate data and generate prioritized lists of entities. While this goal has been fulfilled, other more complex tasks require users to export data (targets, chemicals) to further process them outside of the web application. The present work showcases an example of a post-processing bioinformatics and chemoinformatics pipeline that can be used to further analyze subsets of data obtained from TDR Targets (or any other similar database). The post-processing strategies are implemented in python and are available as a Jupyter notebook [59] for direct use or modification by users.

After post-processing data and generating candidate chemical libraries we went back to the TDR Targets web app to explore resultant hits, and visualize the chemical neighborhood around selected compounds.

Finally, it should be noted that in this study, we explored the trypanocidal activity of just over 5% of the prioritized molecules, suggesting that there is still significant hidden potential in the results of this prioritization.

### New set of compounds with validated activity against *T. cruzi*

From the initial 21 candidates tested in primary assays, eight showed reproducible anti-*T*.*cruzi* activity at fixed concentrations. The top 5 compounds were assessed in a secondary screening, and 4 demonstrated suitable selectivity over Vero cells in dose-response analysis. Figure 5 shows the chemical structures, subgraphs from TDR Targets, and the putative targets for the four prioritized hits. Figure S3 shows the subgraphs for candidate compounds selected for experimental validation.

**Fig 5.**
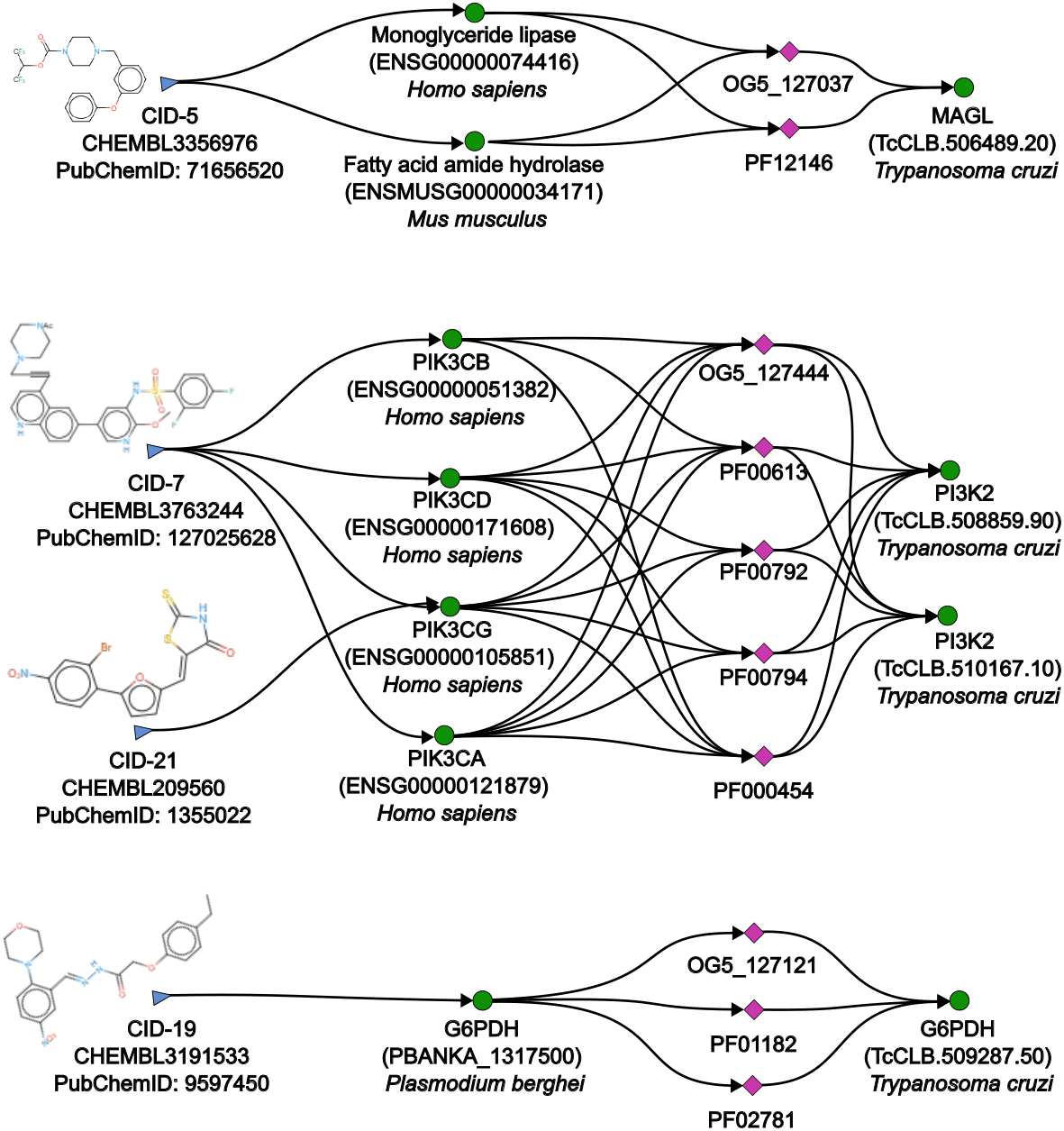
Top 4 hits from the primary screening, extracted subgraphs from TDR Targets, and putative targets. Chemical structures of the top screening hits and the corresponding subgraph retrieved from TDR Targets for each compound. In the graph, blue nodes represent compounds, green nodes represent protein targets, and magenta nodes correspond to functional annotations. The green nodes on the right side of each graph indicate the putative *T. cruzi* targets.

The inclusion of a nitro-containing compound library was chosen due to the validated, though complex, pharmacophore shared by the frontline drugs BNZ and NFX. While this nitro group is associated with potent anti-parasitic activity [43, 60], it can also contribute to off-target reactivity, metabolic toxicity, and genotoxic potential, which are concerns for the existing therapeutics [61, 62]. Among our hits, CID-4 and CID-9 contained nitro groups and displayed moderate cytotoxicity alongside trypanocidal activity, consistent with this challenging profile. However, our pipeline also identified CID-21, a nitro-containing hit, which displays a suitable activity/toxicity profile. This suggests that not all nitro-aromatics share the same promiscuous toxicological profile, and that structural context is critical. Further development of any nitro-containing hit from this library would necessitate rigorous early-stage ADMET profiling to assess whether it offers a genuinely improved therapeutic index over current nitro-heterocyclic drugs.

The decision to experimentally screen a piperazine-based library was predicated on its established role as a privileged, flexible scaffold in medicinal chemistry, valued for its ability to serve as a conformational spacer and present diverse pharmacophores to biological targets. In practice, the piperazine scaffold acts as a molecular dispenser, capable of presenting a diverse array of pharmacophores to biological targets by projecting them into distinct regions of a binding site [63]. The inherent flexibility of the piperazine ring allows it to adopt different conformations [64], meaning that depending on the substituents attached, the molecule can change its conformations (e.g. chair, half-chair, boat, twist-boat), and thus may fit the unique contours of protein pockets [65]. This exploratory, scaffold-driven strategy proved fruitful. Among our hits, the piperazine scaffold was incorporated into chemically distinct architectures, such as in CID-5 and CID-7. Notably, these piperazine-containing hits were computationally associated with different putative target classes (see next section). This divergence supports the core premise of our selection: the piperazine scaffold itself did not confer a single, predetermined mechanism of action but rather facilitated the discovery of structurally novel compounds whose activity likely stems from their unique, appended functional groups. The success of this series validates the inclusion of such ‘framework’ scaffolds in prioritization pipelines, as they can efficiently sample disparate regions of chemical and target space, moving beyond analogues of known inhibitors to uncover genuinely novel chemotypes.

Although one might expect a correlation between library membership and mechanism of action, it is important to note that, at this point, it is impossible to determine whether these scaffolds are actually responsible for any observed trypanocidal activity. Two alternative hypotheses can be proposed about their action: they either inhibit a target or pathway in the host cell that is necessary for the replication of the parasite, or they inhibit a target in the parasite itself. This should be investigated in future work.

Regarding the proposed candidate targets and the observed selectivity (EC_50_ vs CC_50_), it should be considered that we only validated a small number of compounds for practical and economical reasons, and that these were mostly derived from bioactivity evidence from model organisms (human, mouse, rat, etc.). Hence while it provides a proof of concept that the prioritization strategy can yield active compounds, it also serves to highlight the importance of carefully assessing the selectivity of all the candidates, and other analogs when undertaking the experimental validation of the other ∼300 compounds from the 14 remaining libraries produced in this work. As an example it is worth noting that the strategy relies on repositioning of compounds to *T. cruzi* from often complex topologies of the network subgraphs that connect targets, and compounds (number of nodes, multiplicity of interconnections, co-occurrence of positive and negative bioactivity associations, see Figure 5). Hence there is ample room for improving these hit compounds further through exploration of the chemical space around them, and when validating their respective targets and mechanisms of action.

### Putative Targets of the validated hits

Concerning the proposed targets of the 4 prioritized compounds, CID-5 appears as a possible inhibitor of a monoacylglycerol lipase, MAGL (TcCLB.506489.20). Interestingly, CID-7 and CID-21, which belong to the piperazine and nitro libraries, respectively, and showed a low Tanimoto similarity index (0.387), were both associated with the same *T. cruzi* target: phosphatidylinositol 3-kinase 2, PI3K2 (TcCLB.510167.10). Finally, CID-19 and CID-6 both appeared as putative inhibitors of glucose-6-phosphate dehydrogenase, G6PDH (TcCLB.509287.50). All three targets have a high NDS (Figure 6). A brief discussion on the putative targets for primary screening hits not included in the secondary round can be found in the Supplementary Material.

**Fig 6.**
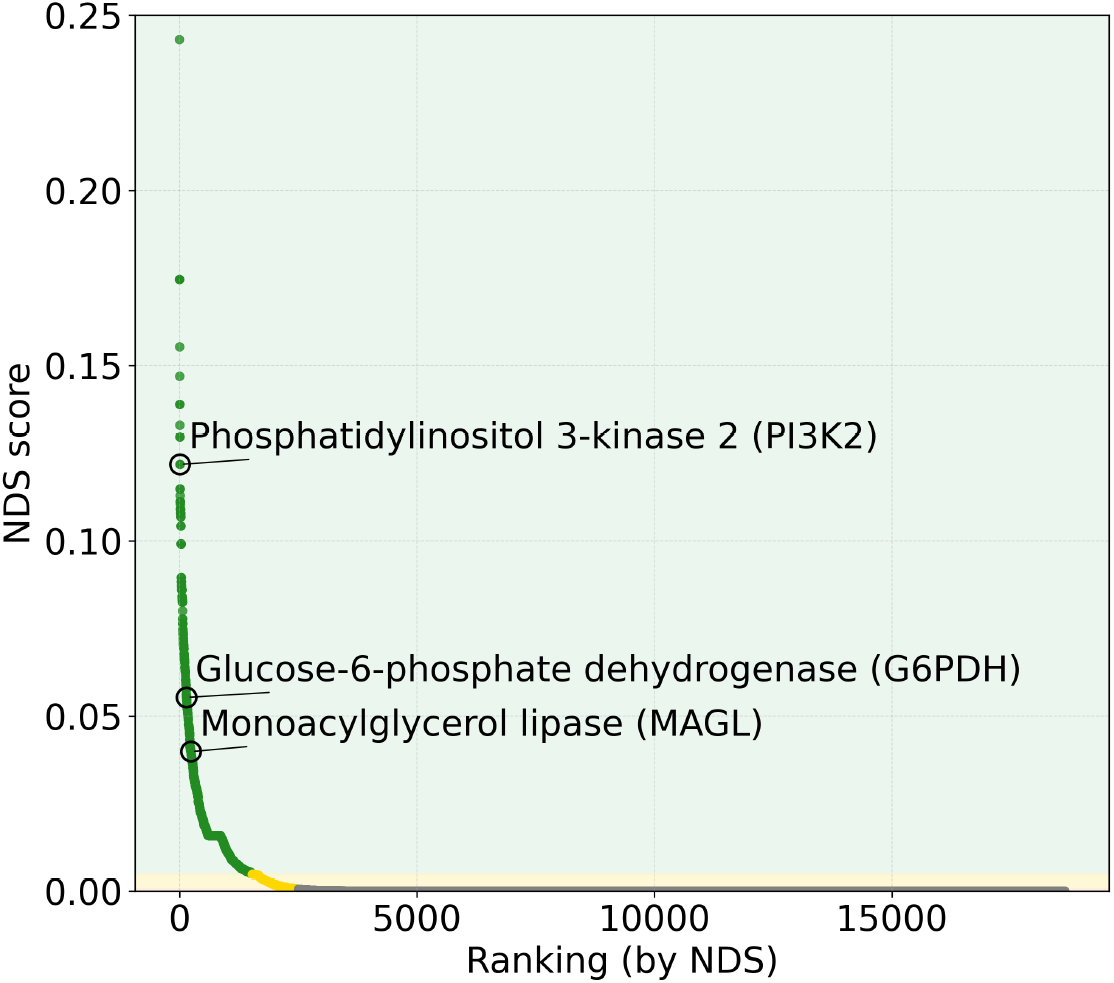
Ranking of the *Trypanosoma cruzi* proteome based on the Network Druggability Scores (NDS). Full ranking of all protein-coding genes in *T. cruzi* based on their NDS. The x-axis represents genes ranked in descending order of their score, while the y-axis shows the quantitative NDS value. This metric reflects the probability of a protein being a successful drug target, derived from its connectivity within a multilayer chemogenomic network and the enrichment of bioactive compounds in its functional neighborhood. High-confidence candidates are located at the far left of the ranking, where scores are highest. The shaded background delimits statistical confidence zones: the green zone (highest confidence - DG 5 and 4), the yellow zone (DG 3), and the red zone (low-confidence targets below baseline noise - DG 2 and 1). The proposed targets for active compounds are identified by labels.

It is important to note that the drug-target associations obtained by the network model in this work are fallible, so it cannot be stated with certainty that the targets of these molecules are unequivocally the ones indicated. In the discussion that follows, we worked under the hypothesis that the proposed targets are a sufficient approximation for making decisions about which of the molecules (and their respective putative targets) are worth continuing to study.

G6PDH is a widely studied target in *T. cruzi* [66, 67]. In addition to participating in the glycolytic pathway for energy generation, it is believed to have a defensive/regulatory role against oxidative stress caused by the environment [68, 69], which positions this enzyme as a possible secondary target for combination therapies with BNZ or NFX.

PI3K2 is a member of a kinase superfamily involved in signal transduction that drives various complex cellular processes, such as cell growth, proliferation, differentiation, motility, and trafficking, among others. In trypanosomatids, this and other kinases from the same superfamily have been evaluated as potential therapeutic targets [70]. In screenings with known human PI3K inhibitors, some compounds like NVP-BEZ235 (Dactolisib - an anticancer drug that inhibits PI3K and mTOR in mammals and inhibits the cell growth of cancer cells) showed sub-nanomolar potency [71], and its effectiveness was demonstrated both *in vitro* and *in vivo* for *L. donovani* and *T. brucei* [72]. This precedent provides fertile ground for the repositioning of human PI3K and mTOR inhibitors. However, all clinical trials associated with this drug have been canceled or halted, with the most successful one reaching Phase 2 due to lack of improved efficacy and poor tolerability [73]. In contrast, the molecules found in this work and potentially associated with this enzyme are not structurally related to each other, or to Dactolisib, or other molecules previously studied in trypanosomatids, thus offering two possible new families of inhibitors for this enzyme.

Finally, monoacylglycerol lipase (MAGL, EC 3.1.1.23) is the enzyme that catalyzes the degradation of monoglycerides (intermediate lipids derived from phospholipid degradation) into glycerol and fatty acids, with arachidonic acid as the main chemical species, due to its abundance but also due to its relevance in various signaling cascades that regulate synaptic function and inflammation in mammals [74]. In trypanosomatids, its function is not clear, but there is evidence that it may act as a virulence factor, possibly modulating the host’s innate immunity [75, 76]. To date, there are no reports of MAGL inhibition in trypanosomatids; CID-5 would be the first reported compound capable of inhibiting this enzyme in *T. cruzi*. All enzymes involved in eicosanoid metabolism are extensively studied in mammals, and there are currently multiple inhibitors for the entire pathway and for MAGL in particular, which presents a rich scenario for both direct drug repositioning and for the biological characterization of the enzyme and its study as a potential therapeutic target.

## Conclusion

As shown in this work, it is possible to create screening libraries using TDR Targets, as it enables not only the prioritization of targets but also the expansion of compound series through both existing and inferred relationships between entities within the network. This means that, for any given target of interest, potential inhibitors can be identified based on evidence obtained for homologous proteins in other organisms, as well as structurally similar compounds to a known inhibitor, based on proximity and relevance calculations within the network.

Here, the data repository was very valuable for the initial generation of datasets filtered by features of interest (such as druggability or novelty), but external resources had to be used to programmatically obtain commercial availability or to gain some intuition about the structural diversity explored in the final set of molecules. Despite these difficulties, a small number of molecules whose trypanocidal activity could be evaluated *in vitro* was obtained. Of the roughly 20 molecules tested, 4 showed micromolar anti-*T. cruzi* activity with a suitable cytotoxicity profile in the final dose-response characterization. Because the 20 molecules were not selected following a rationale (beyond novelty for trypanosomatids, commercial availability, and affordability), it could be argued that the “4/20” ratio (active molecules/total molecules) is significantly higher compared to a blind lead search. It must be noted that this calculation does not constitute a conventional success rate and it can hardly be extended to the entire computational prioritization pipeline, since only a few molecules from two of the 16 generated libraries were chosen for experimental testing. However, it is encouraging that the number and quality of the hits validated here are similar to those obtained from similar computational approaches [77–79].

An additional possible advantage of this type of approach is that the prioritization of molecules comes with their putative target. Although there are no guarantees that the recovered target is exactly the one suggested by TDR Targets, the association is highly probable. This provides an actionable starting point for molecules with an interesting trypanocidal profile, and also an extra piece of information to re-prioritize if it becomes necessary to further narrow down the number of molecules: if the target, for whatever reason, is not interesting, there is no point in proceeding with its validation, much less with the optimization of the molecule itself. In this work, putative inhibitors were obtained for G6PDH, MAGL, and PI3K2. Of these, the MAGL enzyme is possibly the most interesting target that emerged from this study. At the same time, orthologs of this enzyme are extensively studied in model [80–82] and pathogenic [83–86] organisms, which offers a large repertoire of available molecules for repositioning and biochemical assays to adapt.

Throughout this work, the potential of using integrative chemogenomics for drug discovery and repositioning in neglected diseases has been demonstrated, with a special focus on Chagas disease. Our results serve as the foundation for new lines of research focused on the validation of putative targets, the expansion of chemical series based on the identified hits, and the elucidation of the possible mechanisms of action involved in their trypanocidal activity.

## Supporting information

**S1 Fig. Dose-response re-test of selected hits against *Trypanosoma cruzi* and Vero cells**. Scatter plots representing the relation between T. cruzi growth inhibition (%) and Vero cell viability (%) at three different concentrations: (A) 20 *µ*M, (B) 2 *µ*M, and (C) 0.2 *µ*M. Data points represent the mean and standard deviation (SD) of two independent biological replicates. The horizontal dashed red line indicates the cytotoxicity threshold (35% viability), and the vertical dashed red line indicates the anti-parasitic activity threshold (60% inhibition). CN: Negative Control (Culture medium); BNZ: Benznidazole (Positive control); DMSO: Dimethyl sulfoxide (Vehicle control); 2, 3, 7, 15, 21: CID numbers of the re-tested compounds.

**S2 Fig. Drug-target subgraphs for candidate compounds selected for experimental validation**. Schematic representation of the subgraphs obtained for each drug-target pair within the dataset. Molecules acquired for experimental validation are distinctively labeled: compounds from the nitro library are shown in red, while those from the piperazine library are shown in green, each accompanied by its corresponding TDR Targets platform numerical identifier. Additionally, 2D chemical structures are provided to illustrate the structural diversity explored in this study. Compounds marked with a star indicate those that yielded positive results in the primary screening (see experimental validation section). Each gray node represents one or more protein targets or intermediate molecular entities within the interaction network.

**S3 Fig. Results of the** *β***-galactosidase inhibition assays**. Active compounds were evaluated for *β*-galactosidase inhibition using parasite lysates. IPTG was included as a positive control. (A) *β*-Galactosidase activity was measured at a fixed concentration of CID17 (50 *µ*M), CID8 (100 *µ*M), CID12 (100 *µ*M), and CID22 (100 *µ*M). (B) Dose-response curve of IPTG, confirming assay sensitivity. Panels A and B correspond to one experiment performed with three technical replicates. (C-F) Dose-response analyses of CID8 (C), CID12 (D), CID17 (E), and CID22 (F). Each of these panels corresponds to one experiment performed with a single replicate. None of the tested compounds showed inhibitory activity against *β*-galactosidase under the conditions assayed.

**S1 File. Library Preparation Notebook**. A Jupyter interactive Python notebook used for the preparation and refinement of chemical libraries. The notebook includes annotated code and multiple visualizations that document each step of the workflow, from data integration and filtering to library construction. The notebook is publicly available at this Github repository: github.com/trypanosomatics/TDR-screening in the `network-based-library` directory.

**S1 Table Genes prioritized in this study**. List of the genes prioritized in this study, along with their respective *T. cruzi* CL-Brener genome locus identifiers, putative functions, druggability groups (DG), and number of compounds.

**S2 Table Compounds tested in this study**. List of the 21 compounds experimentally tested in this study, along with their respective sources (supplier), identifiers, SMILES strings, IUPAC names and the Classyfire ontology chemical class.

## Acknowledgments

The authors thank Dra. Valeria Tekiel and Bioq. Agustina Chidichimo (IIBIO - UNSAM - CONICET), for help with *T. cruzi* cultures, stocks preservation and parasite passages. We also thank Laboratorios Elea Phoenix for providing Benznidazole; Dr. Frederick Buckner for donating transgenic Tulahuen -gal parasites; Dr. David Wishart (Classyfire) and Chemaxon (Marvin) for generous provision of materials and software.

## Data availability

All software code, libraries and raw data are available in a public Github repository at https://github.com/trypanosomatics/TDR-screening.

