## Supplementary Figures for "Prioritization of chemical scaffolds using the TDR Targets database: an integrative workflow for *Trypanosoma cruzi* drug discovery"

**This PDF file includes:**

Supplementary Code S1

Supplementary Figures S1 to S3

### SUPPLEMENTARY CODE

**Code S1 consists of an interactive Python notebook used for the preparation and refinement of chemical libraries.** The notebook includes annotated code and multiple visualizations that document each step of the workflow, from data integration and filtering to library construction. The notebook is publicly available at: https://github.com/trypanosomatics/TDR-screening/blob/main/network-based-library/libraryPreparation.ipynb

### SUPPLEMENTARY FIGURES

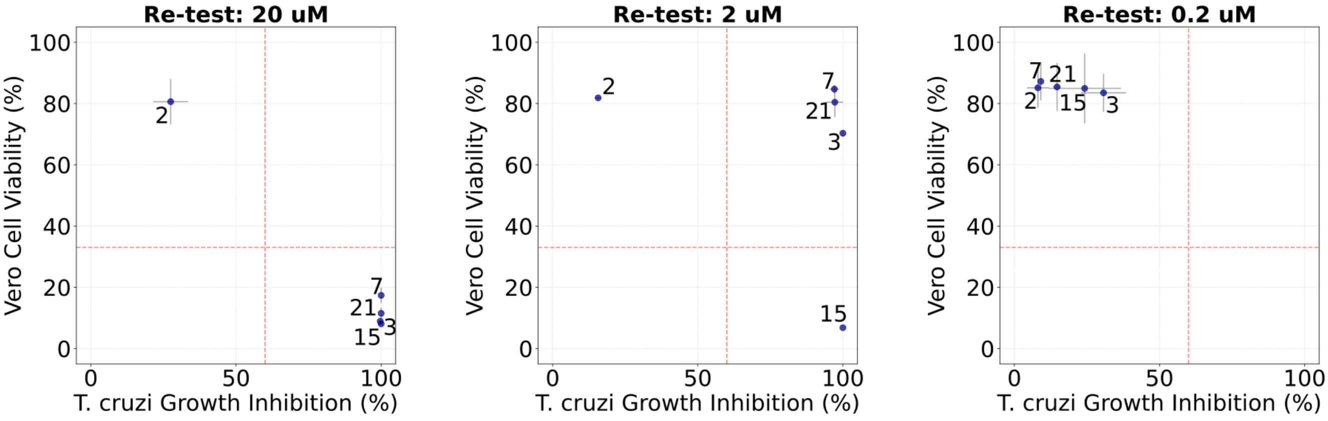


**Figure S1: Dose-response re-test of selected hits against *Trypanosoma cruzi* and Vero cells.** Scatter plots representing the relation between *T. cruzi* growth inhibition (%) and Vero cell viability (%) at three different concentrations: (A) 20 μM, (B) 2 μM, and (C) 0.2 μM. Data points represent the mean and standard deviation (SD) of two independent biological replicates. The horizontal dashed red line indicates the cytotoxicity threshold (33% viability), and the vertical dashed red line indicates the anti-parasitic activity threshold (60% inhibition). CN: Negative Control (Culture medium); BNZ: Benznidazole (Positive control); DMSO: Dimethyl sulfoxide (Vehicle control); 2, 3, 7, 15, 21: CID numbers of the re-tested compounds.

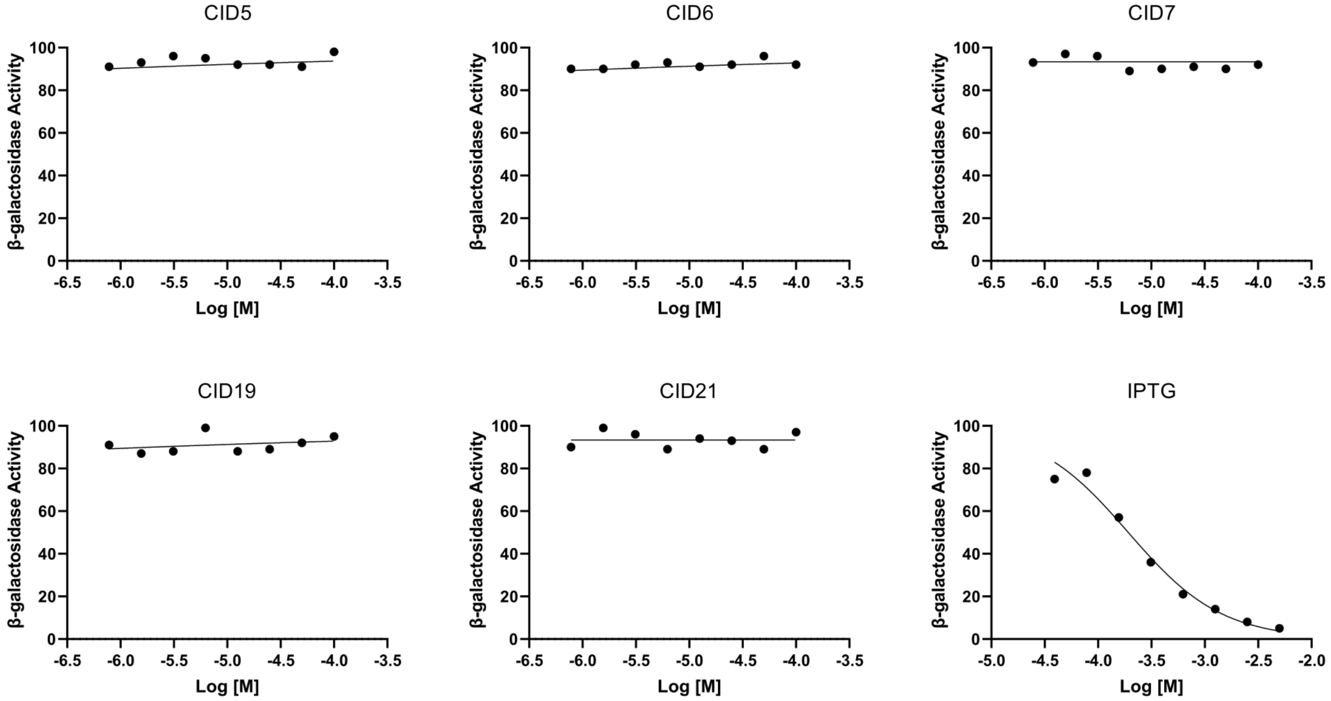


**Figure S2: Results of the β-galactosidase inhibition assays.** Active compounds were tested in a parasite lysate to assess β-galactosidase inhibition assay, using IPTG as a positive inhibition control. None of the compounds displayed inhibitory activity on the reporter enzyme. Each point represents the result of a single biological replicate

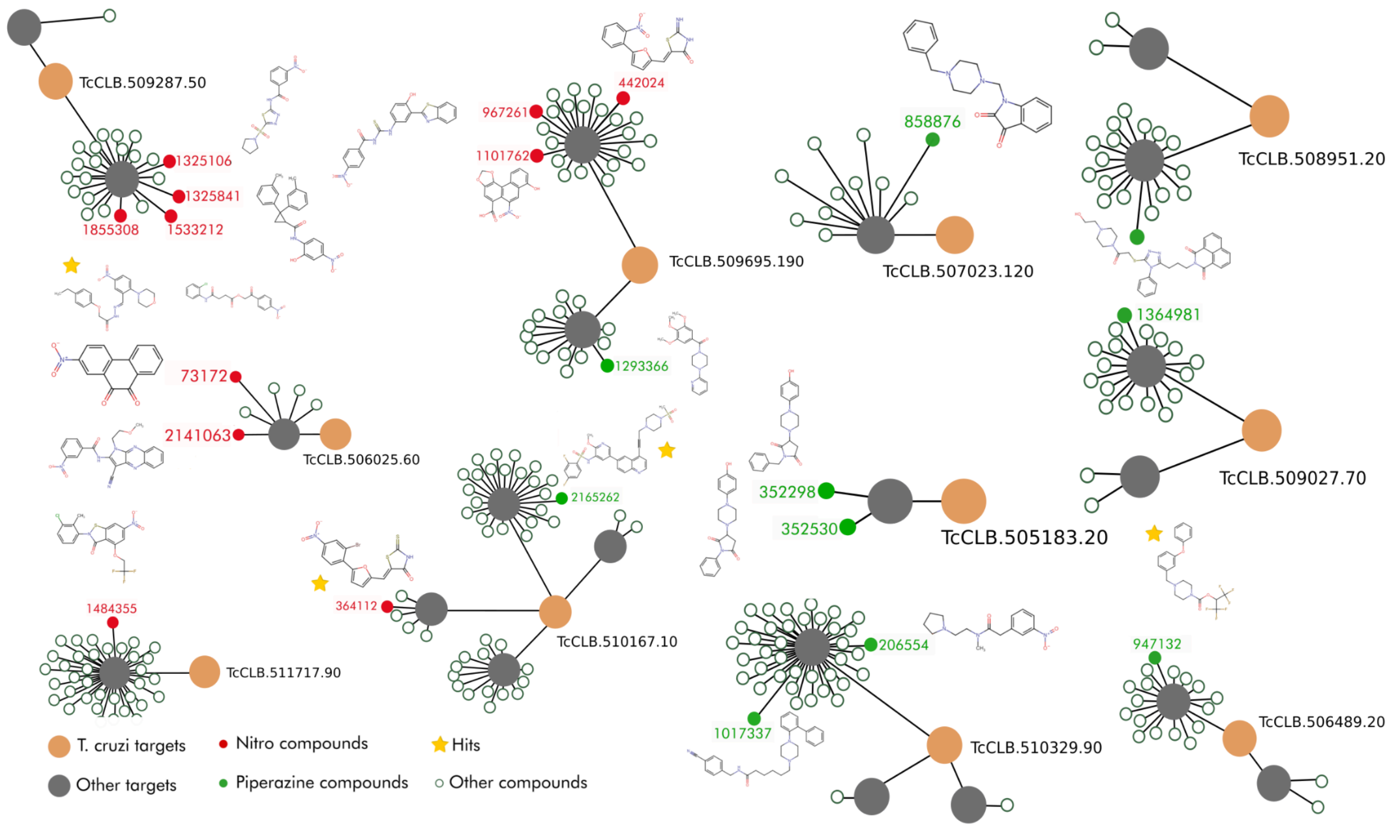


**Figure S3: Drug-target subgraphs for candidate compounds selected for experimental validation.** Schematic representation of the subgraphs obtained for each drug-target pair within the dataset. Molecules acquired for experimental validation are distinctively labeled: compounds from the nitro library are shown in red, while those from the piperazine library are shown in green, each accompanied by its corresponding TDR Targets platform numerical identifier. Additionally, 2D chemical structures are provided to illustrate the structural diversity explored in this study. Compounds marked with a star indicate those that yielded positive results in the primary screening (see experimental validation section). Each gray node represents one or more protein targets or intermediate molecular entities within the interaction network.
